# A comparative study of the human, mouse, and zebrafish eye after staining with RGB trichrome

**DOI:** 10.1101/2022.06.23.497292

**Authors:** Antonio Prieto de la Torre, Francisco Gaytán

## Abstract

Eyesight plays essential roles for the survival of most animal species, and diseases leading to blindness or partial vision loss in humans have an enormous impact in life quality. The complex structure of the eye makes it ideal for the study of structure-function relationships, and to accomplish integration of histology with other disciplines, connecting basic anatomical sciences to clinical medicine. The RGB trichrome consists of the sequential staining with three dyes, fast green (as a general protein stain), picrosirius red and alcian blue (for two main components of the extracellular matrix, collagen and glycosaminoglycans respectively). Notably, this combination of primary colors matches the physiological color detection capacity of the human trichromatic eye. Application of RGB stain to human eye samples and to the eye of two widely used animal models, the mouse and the zebrafish (*Danio rerio*), gives rise to brilliant staining of eye connective tissues, with a range of colors from red to magenta depending on the extracellular matrix composition, and producing highly contrasted tissue interfaces, facilitating the observation of tissue structures, as well as histomorphometric analysis. Interestingly, staining of the retina resulted in differential staining of the retinal layers in the three species analyzed. These results support the effectiveness of RGB as a reliable general staining method, complementary to routine basic hematoxylin and eosin (H&E), applicable to the study of the eye in human pathology, basic preclinical research, as well as in histo(patho)logy teaching.

## Introduction

Vision is critical for most animal survival and constitutes the most important of our senses^**1**^. Accordingly, the largest proportion of the sensorial inputs to our brain comes from the visual system^**2**^. The eye is a complex organ that contains many different histological tissues, the most specific of which is the photosensitive retina, that is a specialized zone of the central nervous system.

Congenital and adquired eye diseases are highly prevalent and result in blindness or partial vision loss that have an enormous impact in life quality. Age-related macular degeneration, glaucoma, cataracts, and diabetic retinopathy are leading causes of vision impairment worldwide^**3-5**^. Due to the relevance of vision, and to the prevalence of specific diseases, the number of studies focused in the eye widely surpasses to those devoted to the rest of sensorial systems^**6**^.

Animal models are essential for basic and preclinical research. The evolution of the eye across species is a fascinating example of the adaptation of an organ to species-specific characteristics^**7**,**8**^. In vertebrate species, the histological architecture of the eye has been highly conserved^**1**^, showing only slight inter-species variations due to adaptation to different lifestyles (i.e., either diurnal or nocturnal) or to different environments (i.e., either terrestrial or aquatic). However, some relevant interspecies differences also exist. Knowledge of the morphological and functional characteristics of the eye in different species, and particularly the similarities and differences with respect to the human eye, is essential for the reliability and translability to the clinic of animal experimental models. The mouse constitutes a classical model for biomedical research, whereas the zebrafish (*Danio rerio*) is an emerging model that is becoming a popular experimental animal for basic research, and both species have been widely used for the study of eye development, physiology, and ophthalmologic disease modeling^**9-14**^.

Histo(patho)logy constitutes the gold standard for the analysis of the structural alterations of eye tissues, either in human pathology or in preclinical studies by using different animal models. Recent advances in image analysis, lead to automatic tissue segmentation, image digitization and virtual histology teaching^**15**,**16**^. These procedures, however, benefit from highly contrasted stained tissue sections, that facilitate the observation and recognition of tissues and structures. Furthermore, in addition to the use of specific immunofluorescence labeling, a general view of the different eye tissues in paraffin embedded samples is suitable. While hematoxylin and eosin (H&E) is the most widely used routine stain for the analysis of tissue structure, there is a wide array of complementary stains for specific tissue components^**17**^. In this context, trichrome stains are used to highlight connective tissue fibers, particularly collagen. The recently developed^**18**^ RGB trichrome (the acronym of the three primary dyes used: picrosirius Red, fast Green, and alcian Blue) combines a collagen specific stain (picrosirius red), a general protein stain (fast green) and a glycosaminoglycan stain (alcian blue).

In this context, it is interesting to know whether the aplication of RGB trichrome, as a basic stain complementary to H&E, is a reliable tool for the study of the human eye, as well as those of the mouse and zebrafish that are two widely used animal models in eye research^**19-21**^.

## Materials and Methods

Human tissue samples were obtained from the Bio-archive of the Department of Pathology of the Faculty of Medicine and Nursing. These specimens corresponded to eyes that were enucleated due to the presence of choroidal melanomas, and were collected between 1980-2000 in keping with contemporary legislation, including informed consent. The use of these samples was approved by responsible members of the Department of Pathology, granting patient confidentiality and following strict adhesion to current legislation.

Animal samples corresponded to mouse and zebrafish eyes. A total of 10 eyes from adult C57BL/6J mice were used, from animals of the vivarium of the University of Cordoba. Zebrafish samples (4 eyes from adult fishes) were obtained from the Servicio de Animales de Experimentación of the University of Córdoba. All animals were euthanized according to the regulations of EU for the protection of animals used for specific experimental purposes. Experimental procedures were approved by the Ethics Commettee of the University of Cordoba.

Human tissues had been fixed in phosphate buffered formaldehyde and embedded in paraffin. A total number of 30 eyes were studied. Areas showing invasion by tumor cells or tissue distortion and/or disorganization due to tumor pressure, were discarded. Animal samples were fixed in Bouin or in phosphate buffered formaldehyde and embedded in paraffin. Six µm thick sections were cut from paraffin blocks, and after dewaxing and rehydration submitted to RGB staining, following previously described methods^**18**^. Briefly, the sections were sequentially stained with 1% (w/v) alcian blue in 3% aqueous acetic acid solution (pH 2.5) for 20 min, washed in tap water, stained with 0.04% (w/v) fast green in distilled water for 20 min, washed in tap water, and finally stained with 0.1% (w/v) sirius red in saturated aqueous solution of picric acid for 30 min, washed in two changes of acidified (1% acetic acid) tap water, dehydrated in 100% ethanol, cleared in xylene, and mounted with synthetic resin. Some human eye sections from the files of the Department of Pathology, that were stained with hematoxylin and eosin or Masson trichrome, were used in order to compare them with RGB-stained tissues.

## Results

The eye is a spherical organ that is made up of three concentric layers that are the retina (the inner photosensitive layer), the uvea (the mid highly pigmented and vascularized layer), and the sclera-cornea (the fibrocollagenous outer coat layer). Many previous studies have described in detail the structure and functional histology of the different components of the eye, and in particular the tissue architecture of the retina, in humans and across species^**1**,**10**,**22-33**^. In this study we will describe the different eye layers in an inside-out direction, the innermost component being the vitreous humor and the outermost component the peri-scleral tissues, and following the usual histological terminology.

The staining characteristics of eye tissues after RGB trichrome application in the three species analyzed were generally similar, although some small differences were observed. We will describe in more detail the RGB staining characteristics of the human eye tissues, and then pointing out the similarities and differences in the mouse and zebrafish tissues with respect to the human samples. A scale bar is shown in one figure per panel, as a reference for comparison. All figures correspond to RGB-stained sections unless otherway indicated in the legend.

### Human eye

All archival human eye samples had been fixed in phosphate-buffered formaldehyde and, therefore, comparisons of the staining characteristics in tissues submitted to different fixation procedures cannot be performed.

#### Retina

The retina is the innermost light-sensitive layer and the most remarkable tissue component of the eye. It contains 5 main neuronal cell types (with different sub-types), that correspond to photoreceptor (cones and rods), bipolar, horizontal, amacrine and ganglion cells, and one main glial cell type (Müller cells). These cell types are organized into three nuclear and two synaptic (plexiform) layers (Fig. 1). Vertical sections stained with hematoxylin and eosin (Fig 1A), classical Masson (Fig. 1B) or RGB (Fig 1C) trichrome allow the observation of these layers. Importantly, in RGB treated sections the main retinal layers appear differentially stained (Figs. 1C,D). So, the outer segments of the photoreceptor layer stain blue whereas the inner segments stain green. This can be observed in either vertical or tangential sections (Fig 2A,B).

**Figure 1.**
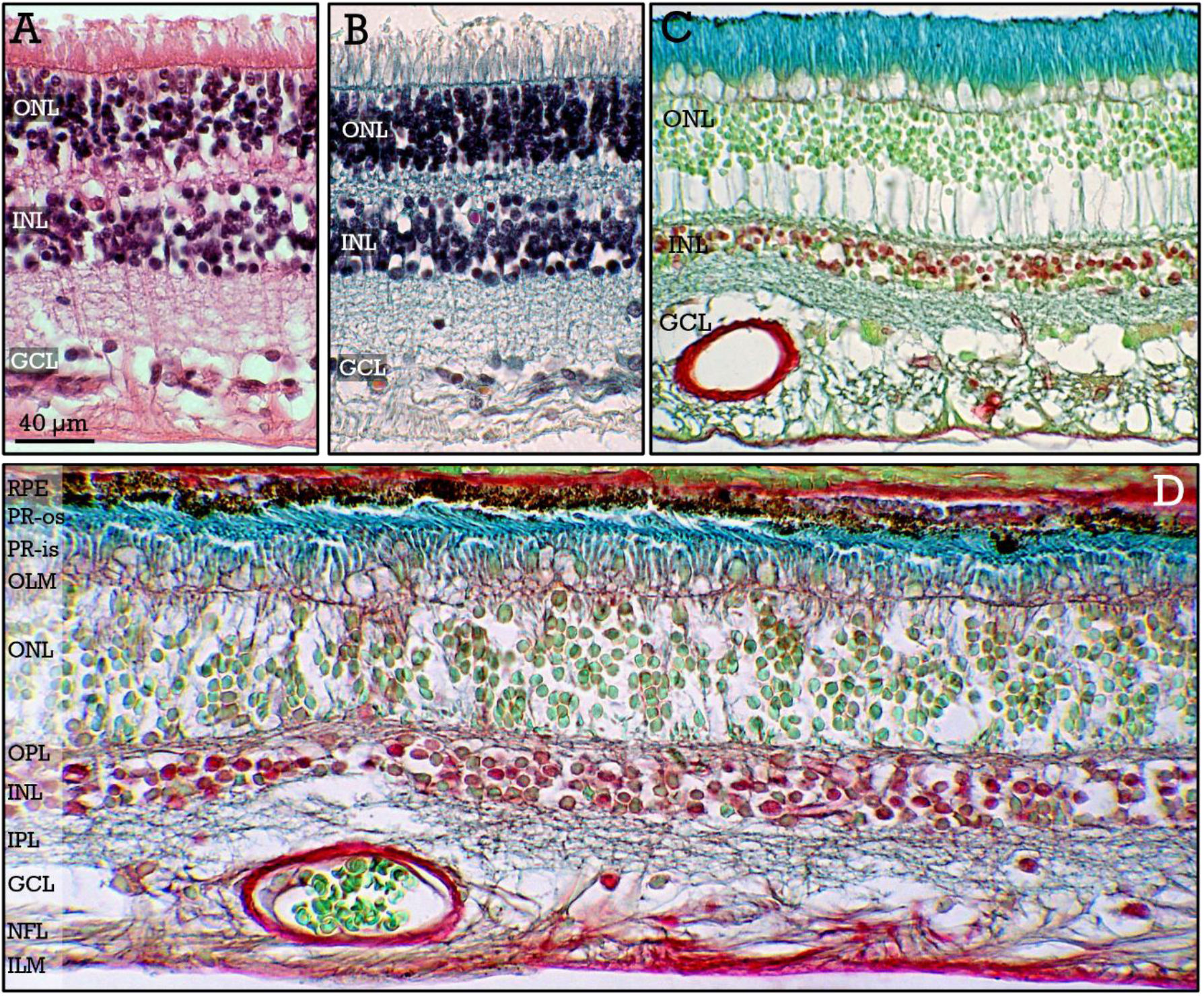
Human retina I. Sensitive peripheral retina stained with H&E (**A**), Masson (**B**) and RGB (**C**) trichrome. The outer (***ONL***) and inner (***INL***) nuclear layers, as well as the ganglion cell layer (***GCL***) are indicated. **D**, vertical section of the retina showing the classical layers: retinal pigment epithelium (***RPE***), photoreceptor outer (***PR-os***) and inner (***PR-is***) segments, outer limiting membrane (***OLM***), outer nuclear layer (***ONL***), outer plexiform layer (***OPL***), inner nuclear layer (***INL***), inner plexiform layer (***IPL***), ganglion cell layer (***GCL***), nerve fiber layer (***NFL***), and inner limiting membrane (***ILM***).

**Figure 2.**
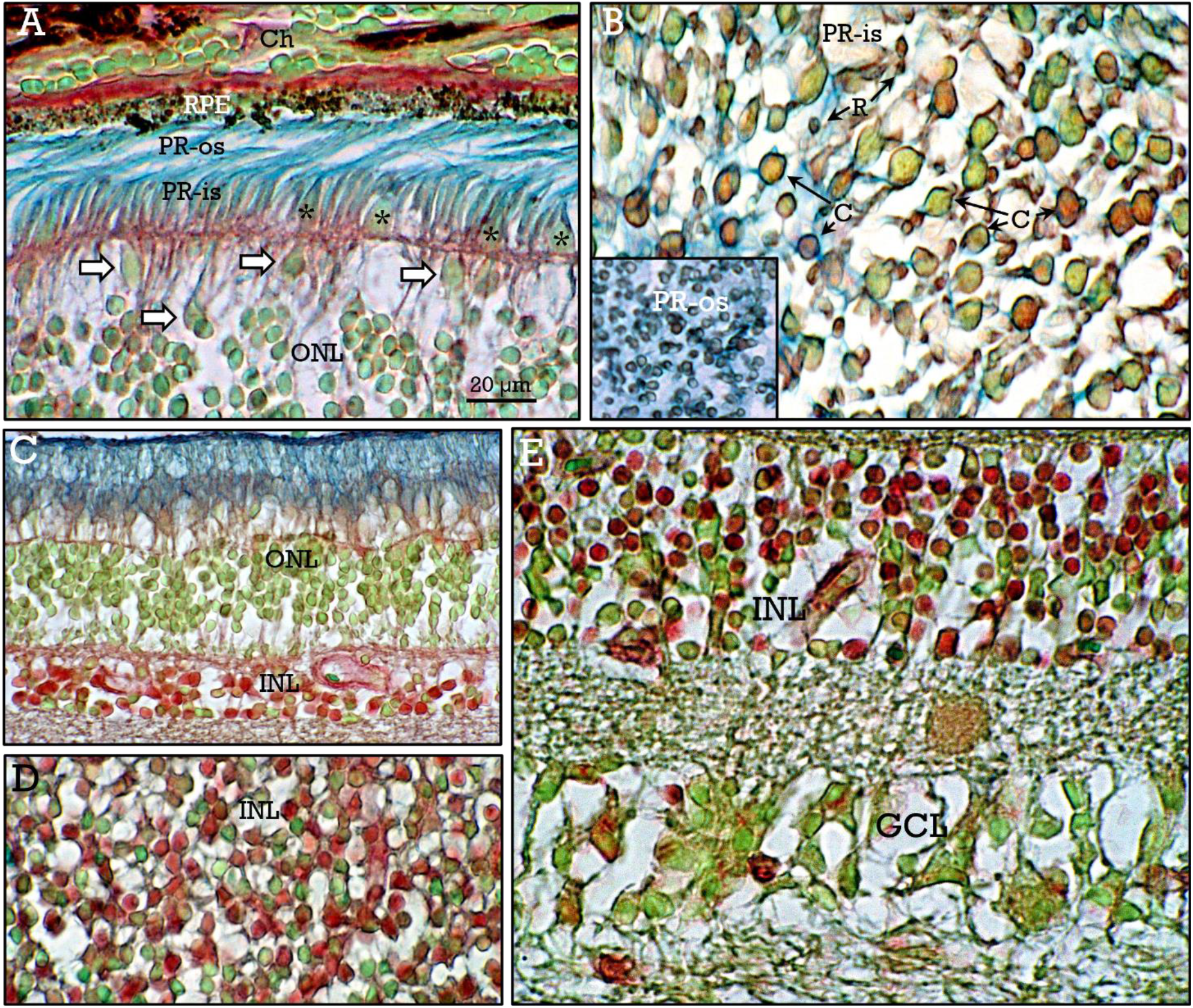
Human retina II. Higher magnification of vertical (**A**,**C**,**E**) and tangential (**B**,**D**) sections of the retina, showing the retinal pigment epithelium (***RPE***) contacting the choroid (***Ch***), the outer and inner segments (***PR-os, PR-is***) of the photoreceptors, and the outer nuclear layer (***ONL***), where some cone nuclei (***white arrows***) can be identified by their elongated shape, their larger size when compared to rod nuclei, as well as their preferential location at the upper zone of the ONL. The inner segments of some cones are labelled by ***asterisks***. A tangential section at the level of the PR-is (**B**) show the inner segments of the cones (***C***) and rods (***R***). In the ***inset*** a tangential section at the level of the PR-is, reveal the blue-stained interphotoreceptor matrix. Differential staining of ONL and INL (**C**,**D**). The heterogeneous staining of the INL nuclei can be appreciated in either vertical (**C**) or tangential (**D**) sections. Some degree of staining heterogeneity can also be apreciated among the cells of the GCL (**E**).

In addition, the outer (ONL) and inner (INL) nuclear layers are also differentially stained with the RGB trichrome (Fig 1C,D). Thereby, in well fixed samples, photoreceptor nuclei in the ONL stain uniformely green. However, at high magnification, the cone nuclei can be distinguised from those of the rods due to their different shape (eliptical vs round), their higher size, and their preferential location at the outermost zone of the ONL (Fig. 2A). In contrast, the INL shows a set of heterogeneously stained cell nuclei, ranging from green to dark red (Figs. 1C,D; 2C,D). The heterogeneity of the staining of INL cell nuclei can be observed in both vertical (Fig. 2C) and tangential sections (Fig. 2D). Heterogeneity in the staining of neurons can be also observed to a lesser extent in the GCL (Fig. 2E).

The classical ten different layers of the sensitive retina, including the outer retinal pigment epithelium (RPE), are indicated in Figure 1D. The nerve fibers of the whole retina leave the globe at the papilla, crossing the *lamina cribosa* of the sclera, becoming myelinated, and giving rise to the optic nerve that connects the retina to the brain (Figs. 3A-C).

**Figure 3.**
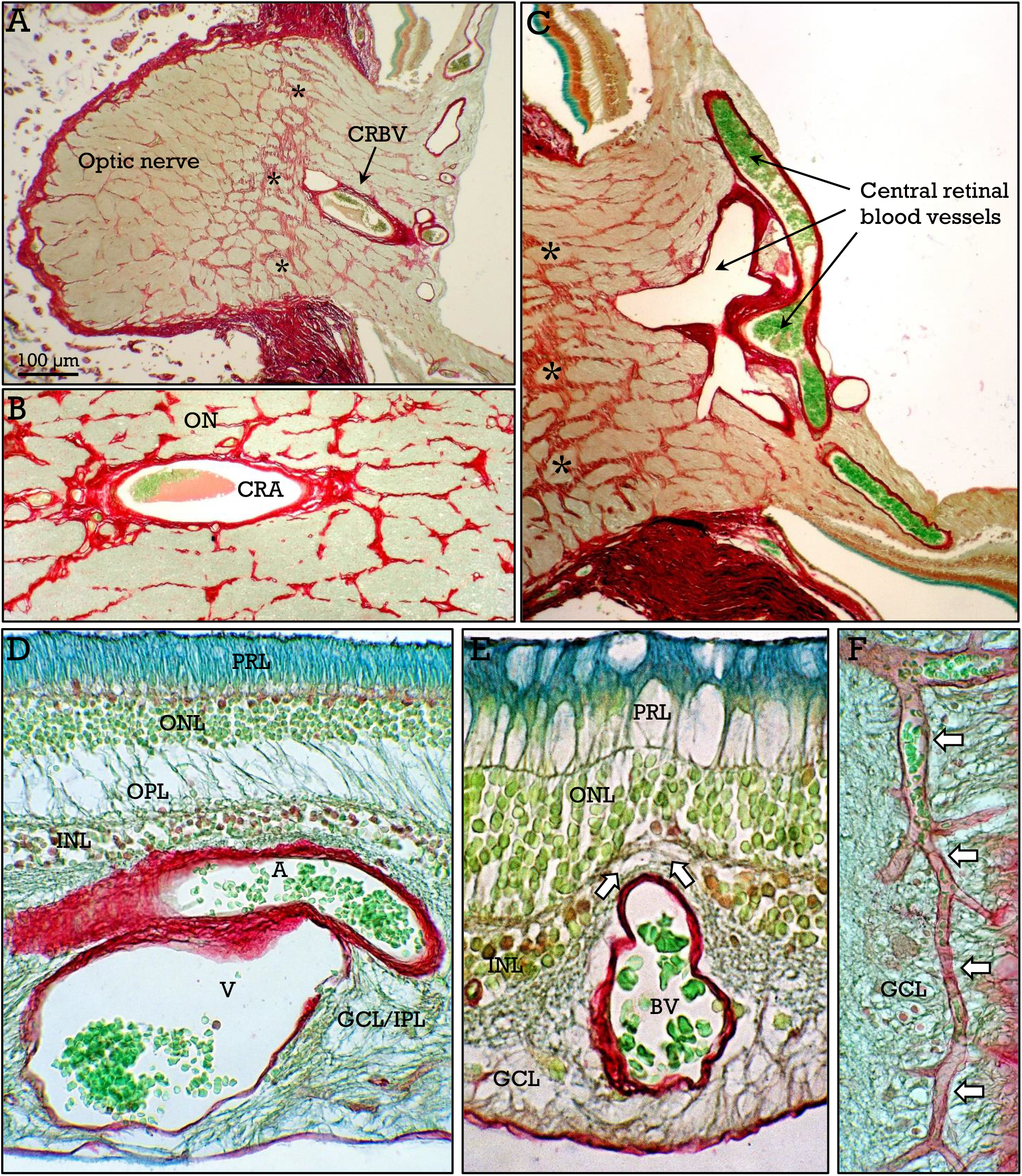
Human retina vascularization I. Central retinal blood vessels (***CRBV***) enter the retina at the point where the optic nerve emerges at the posterior globe aspect (**A-C**). Central retinal artery (***CRA***) runs with the optic nerve (***ON***) and branches once incorporated to the retina. The *lamina cribosa* is marked by ***asterisks***. In **D**, retinal arterioles (***A***) and venules (***V***) run together at the ganglion cells/nerve fiber layers, forming the superficial vascular network. These, relatively large blood vessels (***BV***) displace the INL and even the IPL (**E**), before successive branching into small superficial blood vessels (***arrows*** in **F**).

As a part of the central nervous system, the retina has a high metabolic rate and requires a high supply of oxygen and nutrients, that are provided by a double vascular system. The inner retinal layers (up to the outer plexiform layer) are irrigated by retinal blood vessels, whereas the ONL, the photoreceptor, and the RPE layers are avascular. Blood vessels can be easily identified in RGB stained sections by the red walls and brilliant green-stained erythrocytes (Fig. 3). Retinal blood vessels, derived from the central retinal artery, enter the globe with the optic nerve and branche once they are within the eye (Figs. A-C). Usually, arteries and veins run together (Fig.3D), and these relatively large vessels can displace the inner nuclear and even the outer plexiform layers (Fig. 3E). Large blood vessels at the inner retinal layers, together with those derived from its sucessive branching give rise to the superficial vascular network, located at the ganglion cell/nerve fiber layers (Fig. 3F). Ramification of blood vessels in the superficial network give rise to deep vascular plexus, located below and above the INL (Fig. 4A).

**Figure 4.**
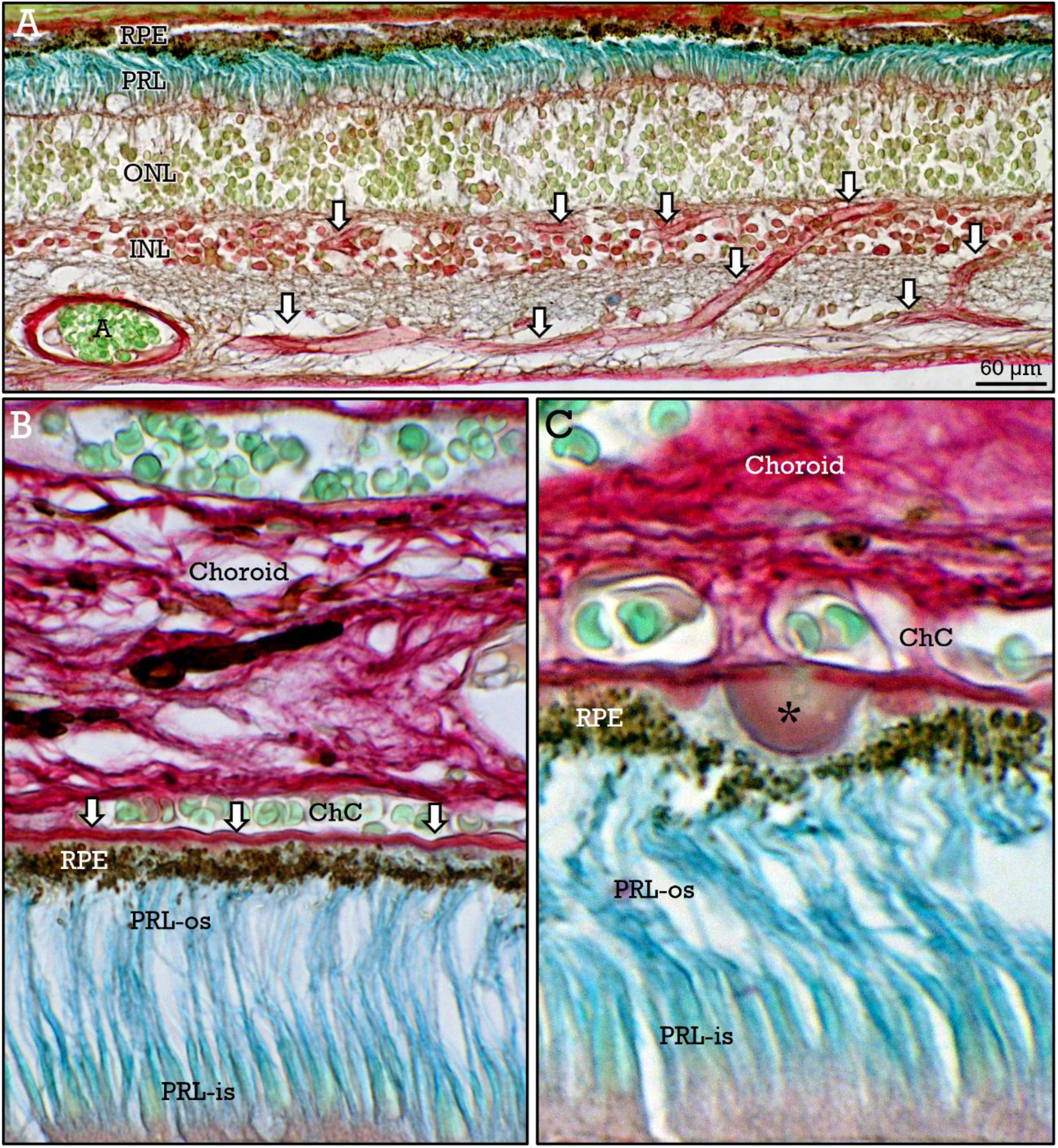
Human retina vascularization II. Ramifications of the blood vessels at the superficial plexus reach the inner plexiforme layer and gives rise to a deep vascular plexus, located below and above the inner nuclear layer (***arrows*** in **A**). Otherwise, outer retina (**B**,**C**) is nourished by choroidal capillaries (***ChC***), adjacent to the Bruch membrane (***arrows***), that separate the choroid from the retinal pigment epithelium (***RPE***). The presence of precipitated material (drusen; ***asterisk***) between the Bruch membrane and the RPE was frequent.

In contrast to the inner vascularized retina, the external retina obtains oxygen and nutrients by diffusion from the adjacent choroidal blood vessels (Fig. 4B). Gases and biomolecules that are interchanged between the outer retina and the choroid have to cross the Bruch membrane, that is located between the RPE and the choriocapillary layer of the choroid (Fig. 4B). Diffusion through this membrane is crucial for retinal pigment epithelium and photoreceptor integrity. So, subretinal deposits of chemically complex material (drusen) under the Bruch membrane (Fig. 4C), distort the RPE and photoreceptor segments, and were frequently observed in samples from elderly patients. The drusen seem to play a role, as an early factor, in age-related macular degeneration^**34**^.

#### Uvea

The uvea constitutes the highly pigmented and vascularized mid layer of the eye (Fig. 5). From behind forward, it is subdivided into the choroid, the ciliary body, and the iris. The choroid lies between the retina and the sclera (Fig. 5A) at the posterior portion of the eye. The red-stained collagen, the brilliant green-stained erithrocytes inside the abundant blood vessels, together with the presence of numerous dendritic brown-stained melanocytes (Figs. 5A-C) clearly mark the limits of this layer. The choroid consists of an inner capillary layer, the choriocapillary, adjacent to the Bruch membrane, a mid layer with small and large blood vessels, and an outer layer of choroid stroma (Figs. 5A,B). Melanocytes are abundant among blood vessels and in the stroma (Fig. 5C). Nerves were also occasionally observed in the choroid (Fig. 5D).

**Figure 5.**
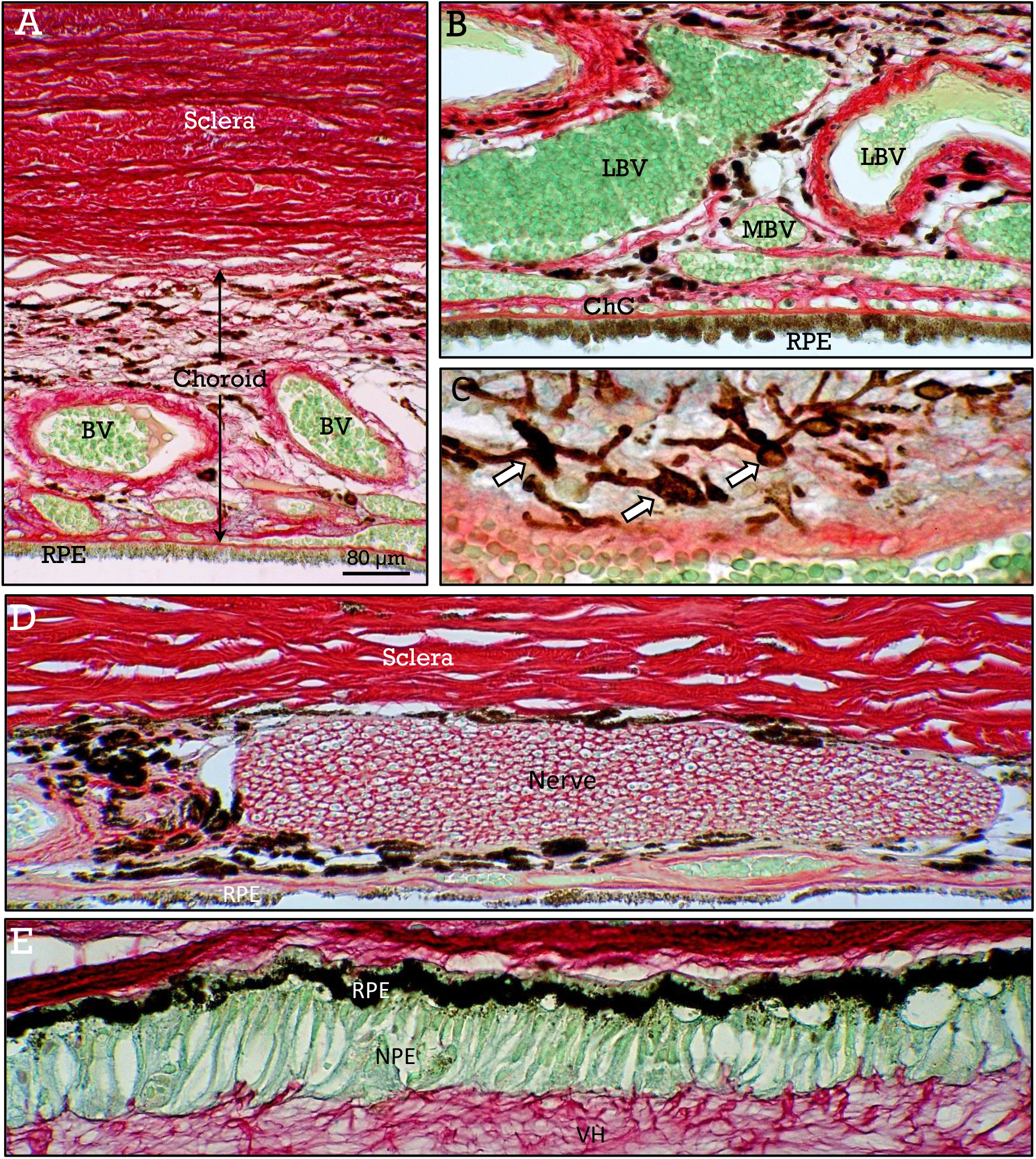
Human choroid. At the posterior aspect, the choroid, located between the retinal pigment epitelium (***RPE***) and the sclera (**A-D**), contains a high number of blood vessels (***BV***). Blood vessels correspond to capillaries at the choriocapillar layer (***ChC***), and to mid (***MBV***) and large (***LBV***) vessels in more external areas. Dendritic melanocytes (***arrows*** in **C**) are very abundant. Occasionally, myelinated nerves were also observed in the choroid (**D**). Anteriorly to the *ora serrata* (**E**) the retina is continous with the *pars plana* of the ciliary body, containing an outer pigmented epithelium (***PE***), and an inner non pigmented columnar epithelum (***NPE***), that is attached to the vitreous humor (***VH***).

Anteriorly to the choroid, the ciliary body have two zones, the *pars plana* and the *pars plicata*. The *pars plana*, that is a non-sensitive extension of the retina anteriorly to the *ora serrata*, is formed by an outer pigmented layer continuous with the retinal pigment epithelium, and an inner non-pigmented layer that is attached to the vitreous humor (Fig. 5E). The *pars plicata* (Fig. 6) is composed of smooth muscle (the ciliary muscle), a highly vascularized stroma, and finger-like projections (ciliary processes). The ciliary projections are covered by two layers of columnar epithelium with an inner layer of non-pigmented secretory cells facing the posterior chamber, and an outer layer of pigmented cells facing the stroma (Fig. 6).

**Figure 6.**
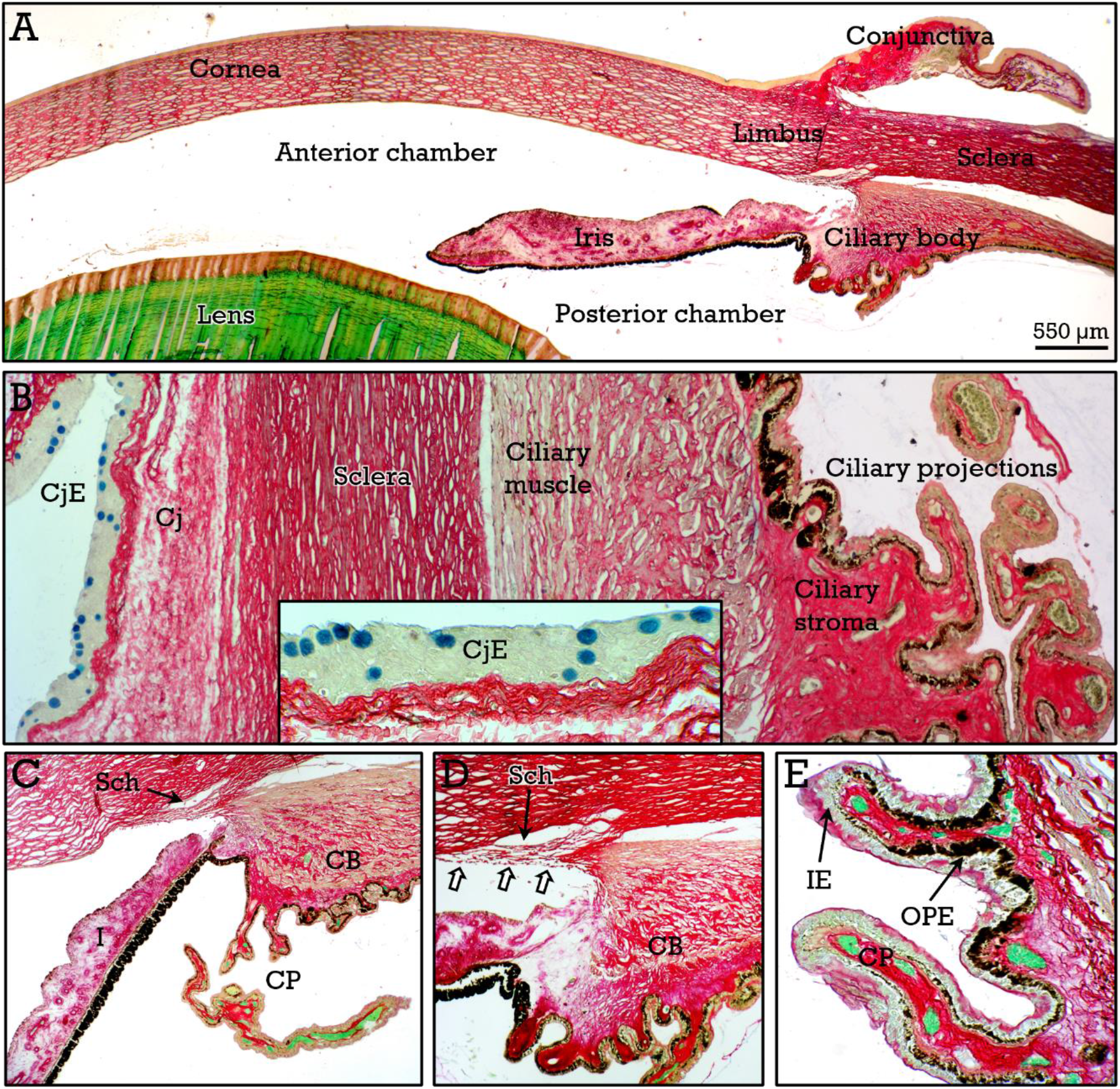
Human ciliary body. In **A**, panoramic view of the anterior zone of the eye, showing the limbus that is the boundary between the sclera/conjunctiva and the cornea, the ciliary body, the lens, and the iris, delimiting the anterior and posterior chambers. In **B**, a panoramic view of the relationships between the ciliary body (***CB***), ciliary projections (***CP***), sclera, and conjunctiva (***Cj***), whose epithelium (***CjE***) contains globet (blue-stained), mucin secreting cells (***inset***). The iridocorneal angle is shown at higher magnification in **C-D**, showing the the trabecular meshwork (***arrows*** in **D**), and the draining Schlemm canal (***Sch***). In **E**, the inner non-pigmented (***IE***) and an outer pigmented (***OPE***) epithelium covering the ciliary projections.

The most anterior portion of the uveal tract is the iris, the colored part of the eye, in the shape of a disc with a central hole (the pupil), that divides the anterior segment of the eye into the anterior and posterior chambers (Figs. 6,7) and is bathed by the aqueous humor. The iris is composed of four layers. These layers are the anterior surface containing stellate fibroblasts and melanocytes, the connective tissue stroma with abundant thick-walled vessels (Figs. 7A-E) and variable numbers of melanocytes (depending on eye color), the iris dilator muscle that appears as a redish line that extents along the iris, anterior to the pigmented epithelium (Fig. 7E). This muscle corresponds to the contractile processes of myoepithelial cells of the posterior epithelium, which is in turn composed of two layers of highly pigmented cells. Furthermore, the iris constrictor (sphincter) muscle (Fig. 7D) appears as a compact bundle of greenish stained smooth muscle cells at the circumpupillary zone. The stroma of the iris appears stained magenta in contrast to the red stained stroma of the ciliary processes (Figs. 7A,B).

**Figure 7.**
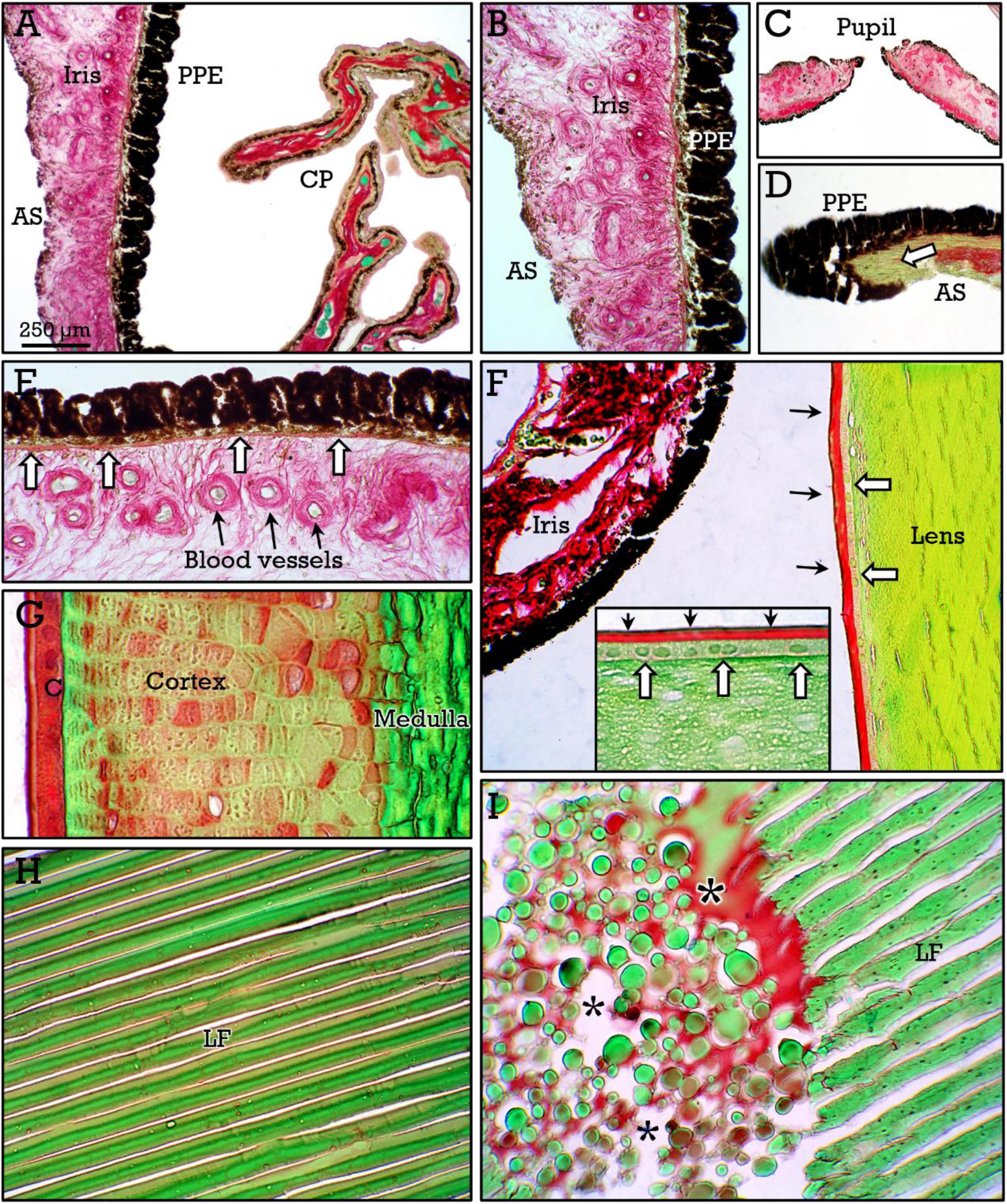
Human iris and lens. Figures **A-E** show the structure of the human iris, showing the anterior surface (***As***), the posterior pigmented epithelium (***PPE***), the pupillae constrictor muscle (***arrow*** in **D**), the pupillae dilator muscle (***arrows*** in **E**), and the stroma containing thick-walled blood vessels (**E**). Figures **F-J** show the structure and staining properties of the lens. The red-stained capsule (***black arrows*** in **F**) contrast with the green-stained mature fibers (**F**,**H**). In **C**, the cortex showing redish staining (**G**). In the anterior face, the lens epithelium (***white arrows*** in **F**) can be observed. In some samples, morgagnian cataracts with liquefaction of lens fibers (***asterisks*** in **I**) were observed.

The structure of the functionally relevant irido-corneal angle, containing the trabecular meshwork and the Schlemm canal, is shown in Figure 6. These structures are responsible for the draining of aqueous humor, that is secreted by the ciliary body epithelium and flows from the posterior chamber into the anterior chamber by passing through the pupil.

#### Crystalline Lens

Lens is attached to the ciliar projections by ciliary fibers (lens zonule) that are hair-like filaments, that are poorly stained and are difficult to observe under light microscopy. Lens have a red-stained capsule (Fig. 7F,G) and a redish-stained cortex containing non-fully mature fibers (Fig. 7G). Toward the center of the lens (medulla), the fibers become homogeneous and dense and appear as green-stained parallel anucleated fibers (Fig. 7H). On the anterior face, the lens surface epithelium can be observed (fig 7F) as a simple layer of cuboidal cells under the lens capsule. In some samples belonging to elderly patients, cataracts were frequently observed (Fig. 7I).

#### Sclera-cornea

The sclera is the outer fibrocollagenous coat of the eye, responsive for the maintenance, together with the humor pressure, of the eye globe shape. At the posterior pole, the sclera forms a sieve-like connective tissue, the *lamina cribosa* of the sclera (Figs. 3A,C) and is continuous with the optic nerve sheath, that is formed by the same membranes that enclose the brain and the spinal cord (Fig. 8A).

**Figure 8.**
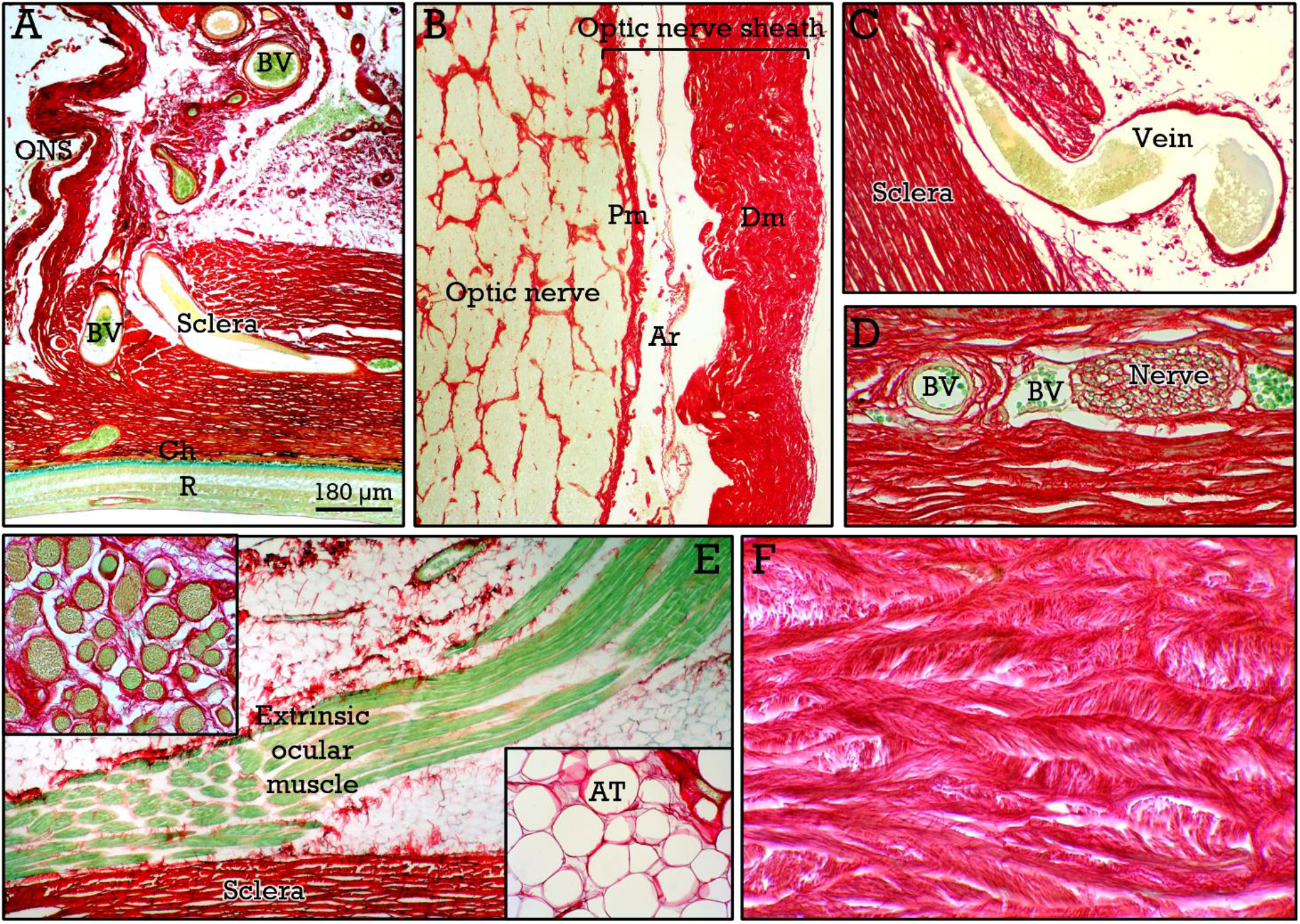
Human sclera. At the level of the optic nerve (**A**,**B**), the sclera is continuous with the optic nerve sheath composed of the dura mater (***Dm***), the arachnoids (***Ar***) and the pia mater (***Pm***). At the outer sclera (**C-E**), blood vessels (***BV***), nerves and the insertion of the extrinsic ocular muscles (***EM***), cut either longitudinally (**E**) or tansversally (***upper inset*** in **E**), together with periscleral adipose tissue (lower ***inset*** in ***E***) can be observed. The dense connective tissue of the sclera is composed of cross-linked collagen fascicles (**F**).

This sheath consists of three covering layers, the pia mater, the arachnoid and the dura mater (Fig. 8B). Although the sclera is poorly vascularized, blood vessels, as well as nerves, are present at the outermost zone (Fig. 8C,D). Extrinsic ocular muscles insert at the outer surface of the sclera (Fig. 8E), and periorbital adipose tissue is also present (Fig. 8E). The scleral stroma is composed of thick collagen bundles, that are oriented in different directions parallel to the surface, forming a cross-linked red-stained mesh (Fig. 8F).

The anterior portion of the sclera is covered by a mucous membrane, the conjunctiva, composed of a stratified epithelium containing globet cells (Fig. 6B) secreting mucins that contribute to the thin fluid layer (tear film) covering the ocular surface. At the limbus, the sclera and the covering conjunctiva are replaced by the cornea (Fig. 9A), that is one of the refractive elements of the eye, together with the crystalline lens. The structure of the cornea (from front to back) consists of the anterior stratified non keratinized corneal epithelium, the Bowman membrane, the corneal stroma, the Descemet membrane and the thin posterior endothelium (Figs. 9B-D). Blood vessels are absent in the corneal stroma. Small areas of the cornea lacking the Bowman membrane and occupied by expanded anterior epithelium (likely due to corneal erosion and related to the lack of regenerative capacity of this membrane) were occasionally observed (Figs. 9E,F) in some samples.

**Figure 9.**
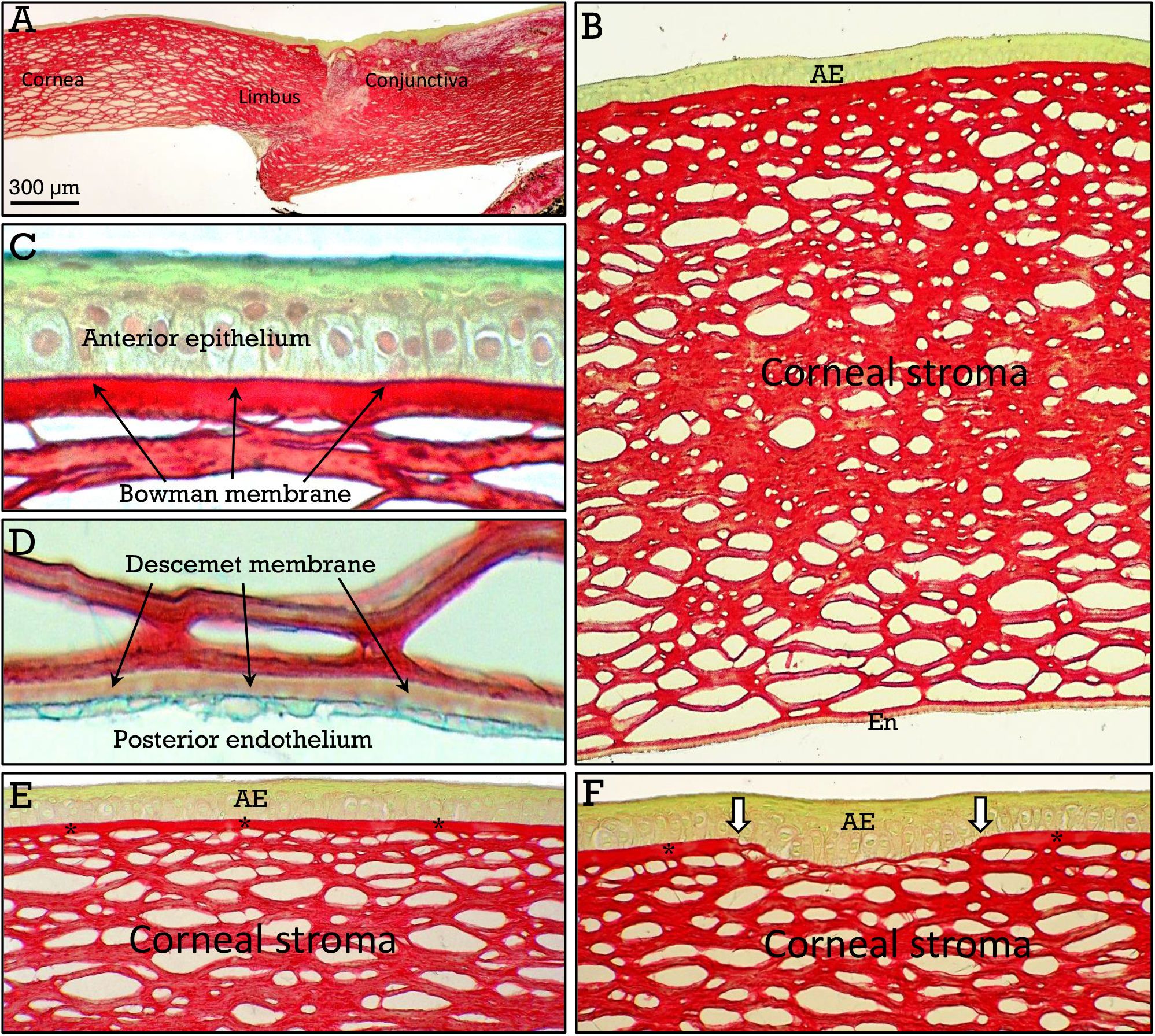
Human cornea. The limbus (**A**) corresponds to the transition from the opaque sclera/conjunctiva to the transparent cornea. The cornea (**B-F**) is composed of an anterior, non keatinized stratified epithelium (***AE***), a collagenous stroma and a posterior endothelium (***En***). The Bowman membrane is located at the basis of the anterior epithelium, whereas the Descemet membrane is anterior to the endothelium (**C**,**D**). The Bowman membrane (***asterisks*** in **E**) was occasionally absent in some small areas (***arrows*** in **F**).

### Mouse eye

Whole mouse and zebrafish eyes were fixed in either 4% phosphate-buffered formaldehyde or Bouin fixative, and the staining characteristics of the RGB staining in samples submitted to different fixation procedures were analyzed.

#### Retina

The general structure of the mouse retina was identical to that of humans (Figs. A-C). With respect to the influence of the fixation process, some differences in RGB staining were observed between Bouin- (Fig. 10A) and formaldehyde- (Fig. 10B) fixed samples. Slightly different shades of colors were found with respect to the uniformely stained ONL and photoreceptor layers (Figs. 10A,B). However, the most relevant difference corresponded to the staining of the INL, that was uniformely stained in Bouin fixed samples, but heterogeneously stained in those fixed with formaldehyde (Figs. 10B, D,E). Similar to that found in the formaldehyde fixed human retina, this heterogeneous staining pattern was found in both vertical and tangential (Figs. 10A,D,E) sections. Tangential sections at the level of the retinal pigmented epithelium reveal the polygonal shape of these cells (Fig. 10F).

**Figure 10.**
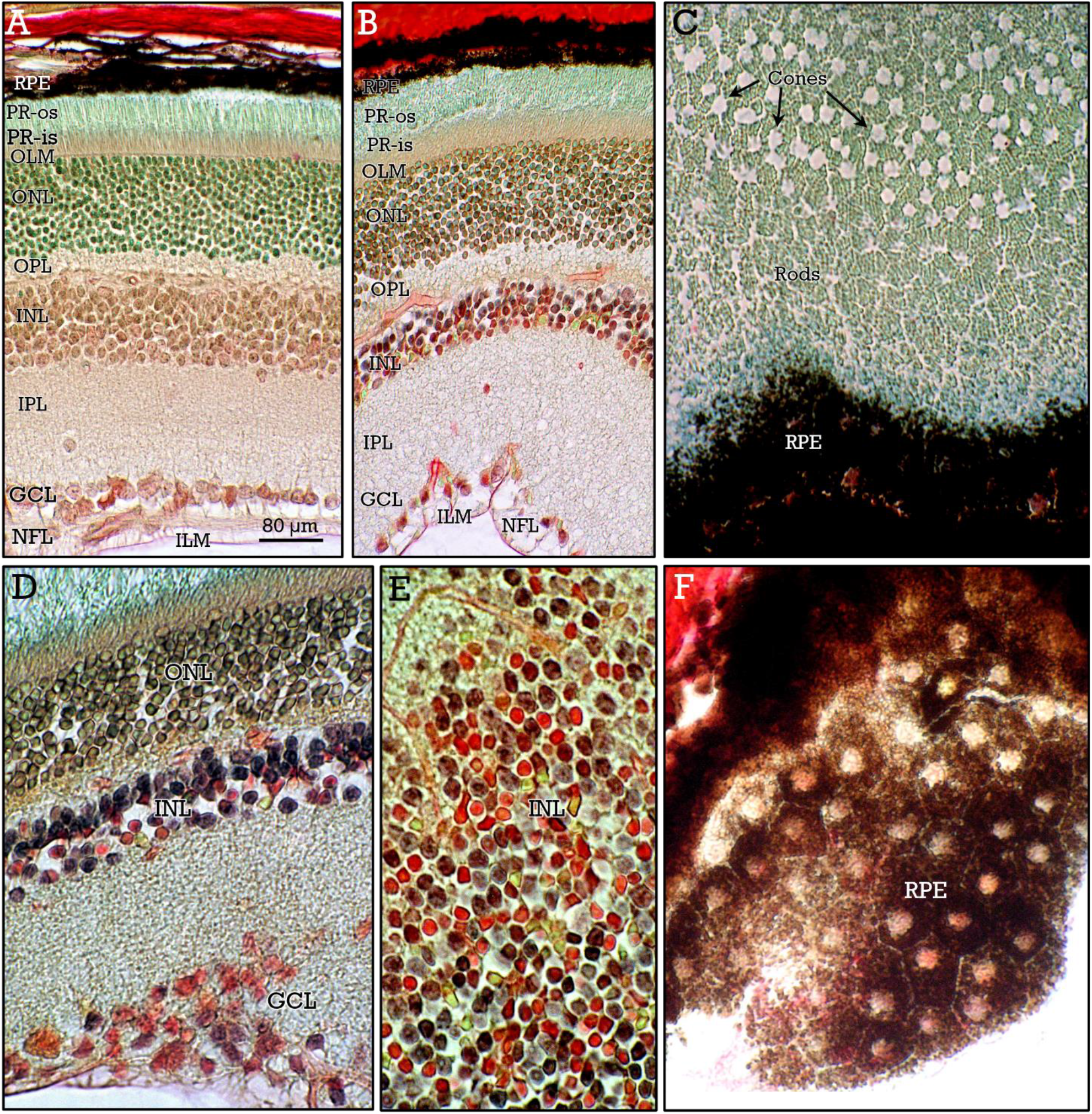
Mouse retina. Vertical sections of mouse retina fixed with Bouin (**A**) or formaldehyde (**B**). The different layers are indicated. Except for subtle differences in the shades of color, the most relevant difference between samples with both fixatives was the heterogeneous staining of the INL and to a lesser extent in the GCL (**D**) in formaldehyde-fixed tissue, similar to that of the human retina. Tangential sections at the level of the photoreceptor layer (**C**), close to the RPE, reveal the cone and rod mosaic pattern. Tangential sections at the level of the INL (**E**) show the heterogeneous staining of the nuclei, while the polygonal shape of RPE cells can be apreciated in tangential sections (**F**).

The patterns of vascularization of the mose and human retina were equivalent. Central retinal blood vessels entering at the papilla with the optic nerve (Fig. 11 A), give rise to superficial (i.e., internal), as well as deep (i.e., external) vascular plexuses at the GCL (Fig. 11B) and the INL (Figs. 11B-D) respectively. In addition, choroid blood vessels were responsible for the supply of oxygen and nutrients to the outer retina layers (Figs. 12 B-D).

**Figure 11.**
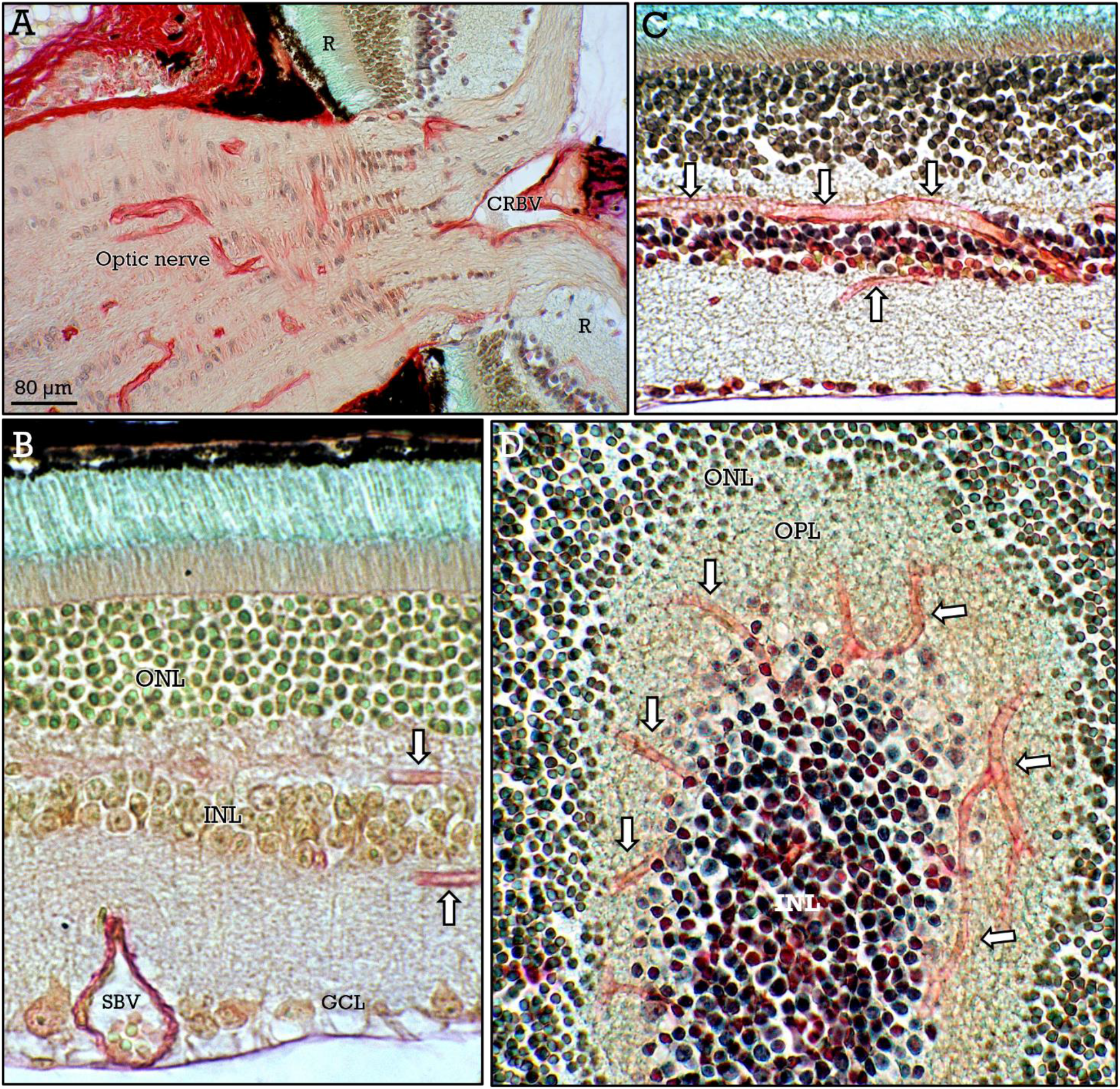
Mouse retina vascularization. Central retinal blood vessels (***CRBV***) enter the retina (***R***) along with the optic nerve. Superficial blood vessels (***SBV***) branch out from the ganglion cell layer(**B**) and form the deep plexus (**B**,**C**) surrounding the INL (***arrows***). The deep plexus (***arrows***) can be clearly appreciated in tangential sections at the level of the INL (**D**).

**Figure 12.**
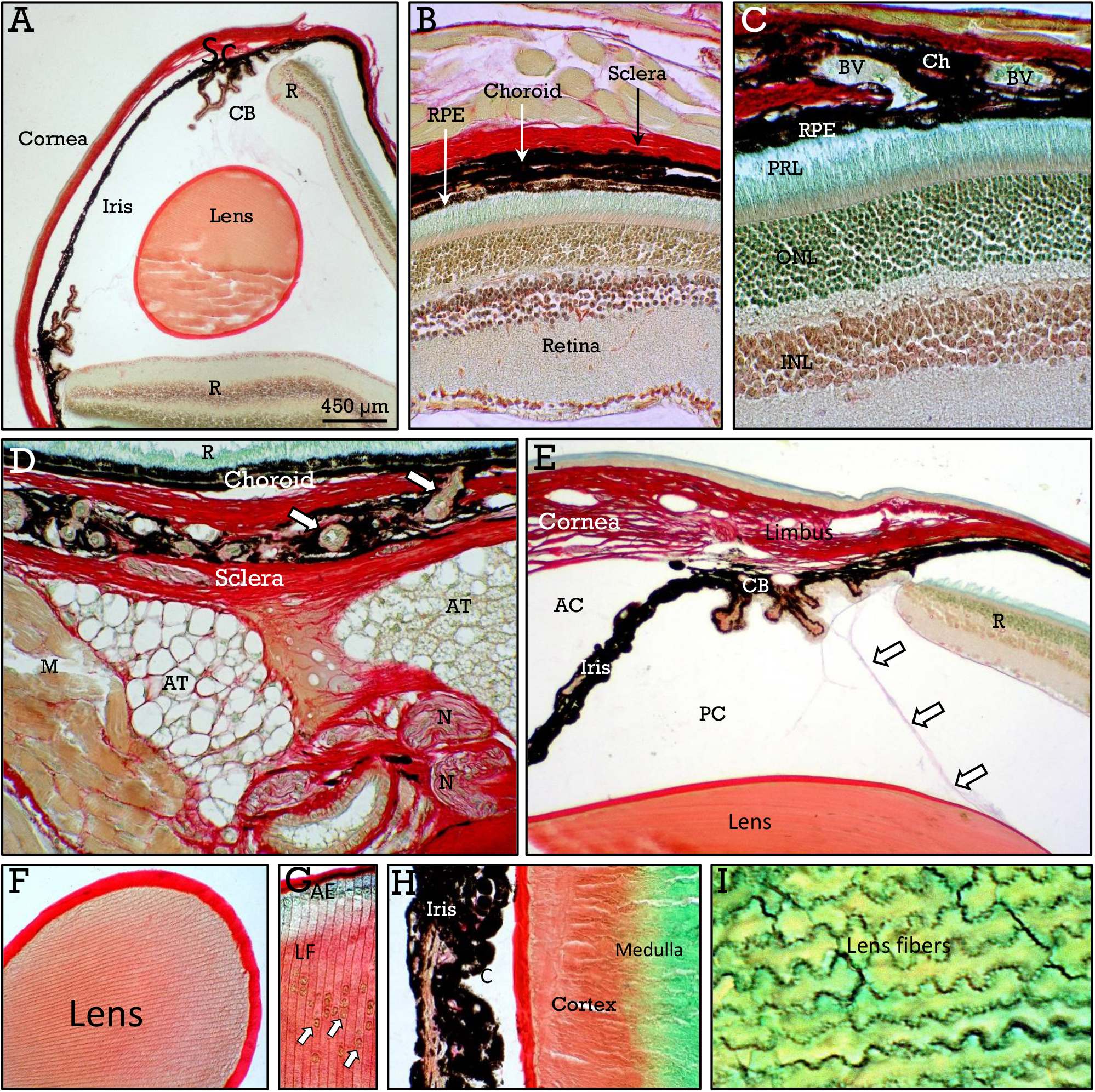
Mouse uvea and lens. A general view of the mouse eye (**A**), shows the retina (***R***), sclera (***Sc***), ciliary body (***CB***), cornea, iris, and lens. **B**,**C**, vertical sections showing the choroid as a thin layer between the RPE and the sclera, containing blood vessels (***BV***), and abundant melanocytes. **D**, choroidal blood vessels (***arrows***) are abundant in the area adjacent to the optic nerve, together with periorbital adipose tissue (***AT***) muscles (***M***) and nerves (***N***). The relationships among ciliary body (***CB***), zonula (***arrows***), lens and the transition between the sclera and the cornea at the limbus, can be appreciated in **E**. Different zones of the lens are shown in **F-I**. The capsule (***C***), cortex and medulla show the same staining properties than human lens. The anterior epithelium (***AE***) and the nuclei of immature fibers at the cortex (***arrows***) are indicated in **G**.

#### Uvea

The mouse uvea, and particularly the choroid, is highly pigmented and difficult to delineate from the retinal pigment epithelium (Figs. 12B,C). Choroidal blood vessels enter the choroid at the peripapillar zone (Figs. 12C,D) and are responsible for irrigation of the outer retinal layers. The ciliary body and the iris, showed a similar, though more pigmented, appearance than their human counterpart (Fig. 12E). The mouse lens also showed the same structure and staining properties than human lens (Figs. 12 F-I).

#### Sclera and cornea

Different tissular structures such as the extrinsic ocular muscles, nerves, blood vessels and adipose tissue can be observed at the outer face of the posterior sclera (Figs. 13A-C). These tissues can be easily distinguised by their morphology and differential staining. At the back area, the sclera is continous with the covering sheath of the optic nerve (Fig. 13 D). As in human eye, the sclera is composed of collagen fascicles oriented in different directions (Fig. 13E). At the anterior zone, the sclera is covered by the conjunctiva and form a underdeveloped nictitating membrane that contains a central lamina of hyaline cartilage (Figs. 14A,B). At the limbus, the sclera/conjunctiva is replaced by the the cornea (Figs. 12A,E), that is made up of the anterior stratified epithelium, the corneal stroma and the inner endothelium (Fig. 14C). Notably, in the corneal stroma, the anterior zone is red-stained whereas the posterior zone is stained magenta.

**Figure 13.**
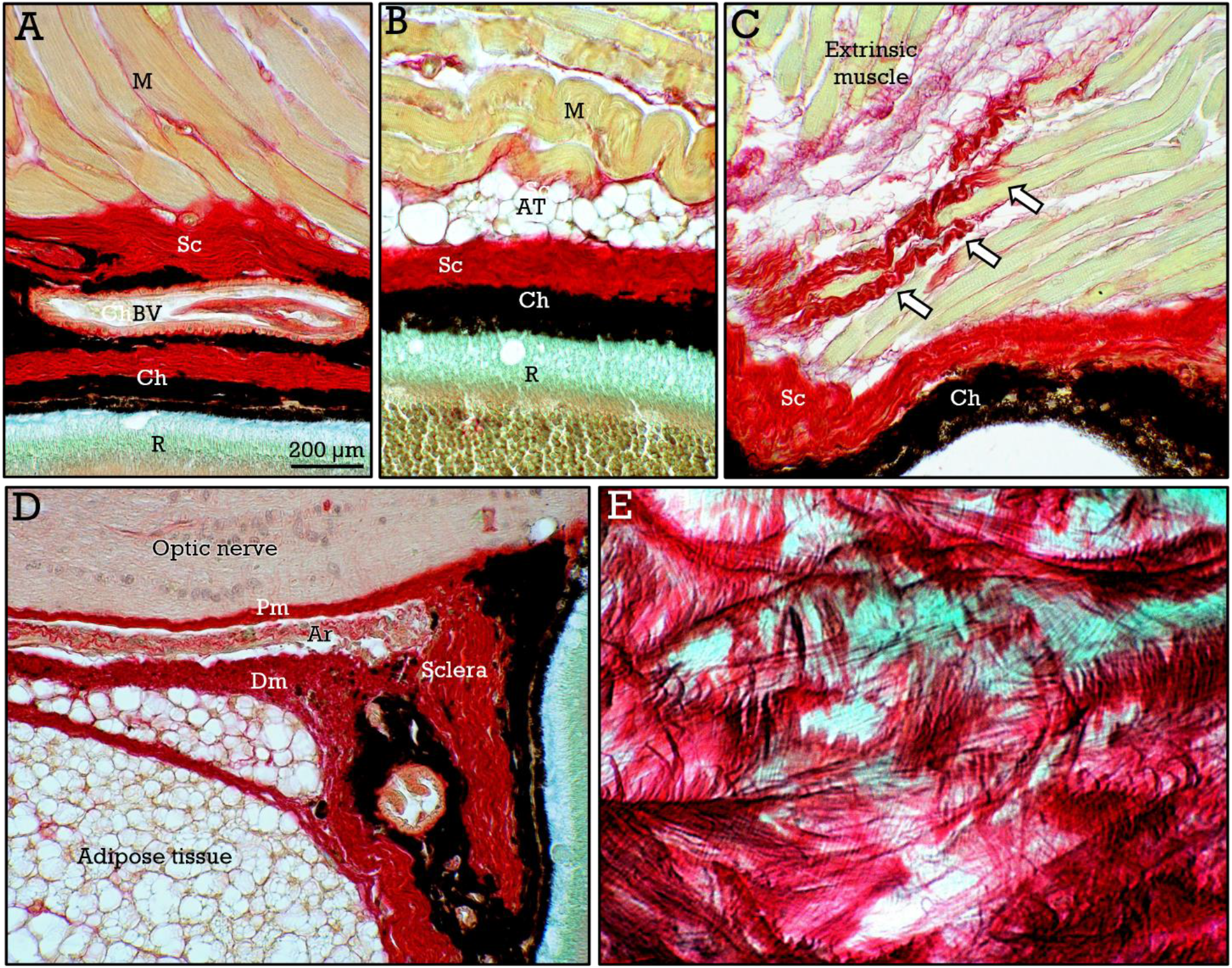
Mouse sclera. The sclera (***Sc***) appears a a red-stained lamina, surrounded by extrinsic ocular muscles (**A**,**B**), and adipose tissue (**B**). Myofibrils of the extrinsic muscles (***EM***) insert in the sclera either directly or through small tendons (***arrows*** in **C**). The sheath of the optic nerve, composed of pia mater (***Pm***), arachnoid (***Ar***) and dura mater (***Dm***), is continuous with the sclera (**D**). Tangential sections show the cross-linked collagen fascicles (**F**).

**Figure 14.**
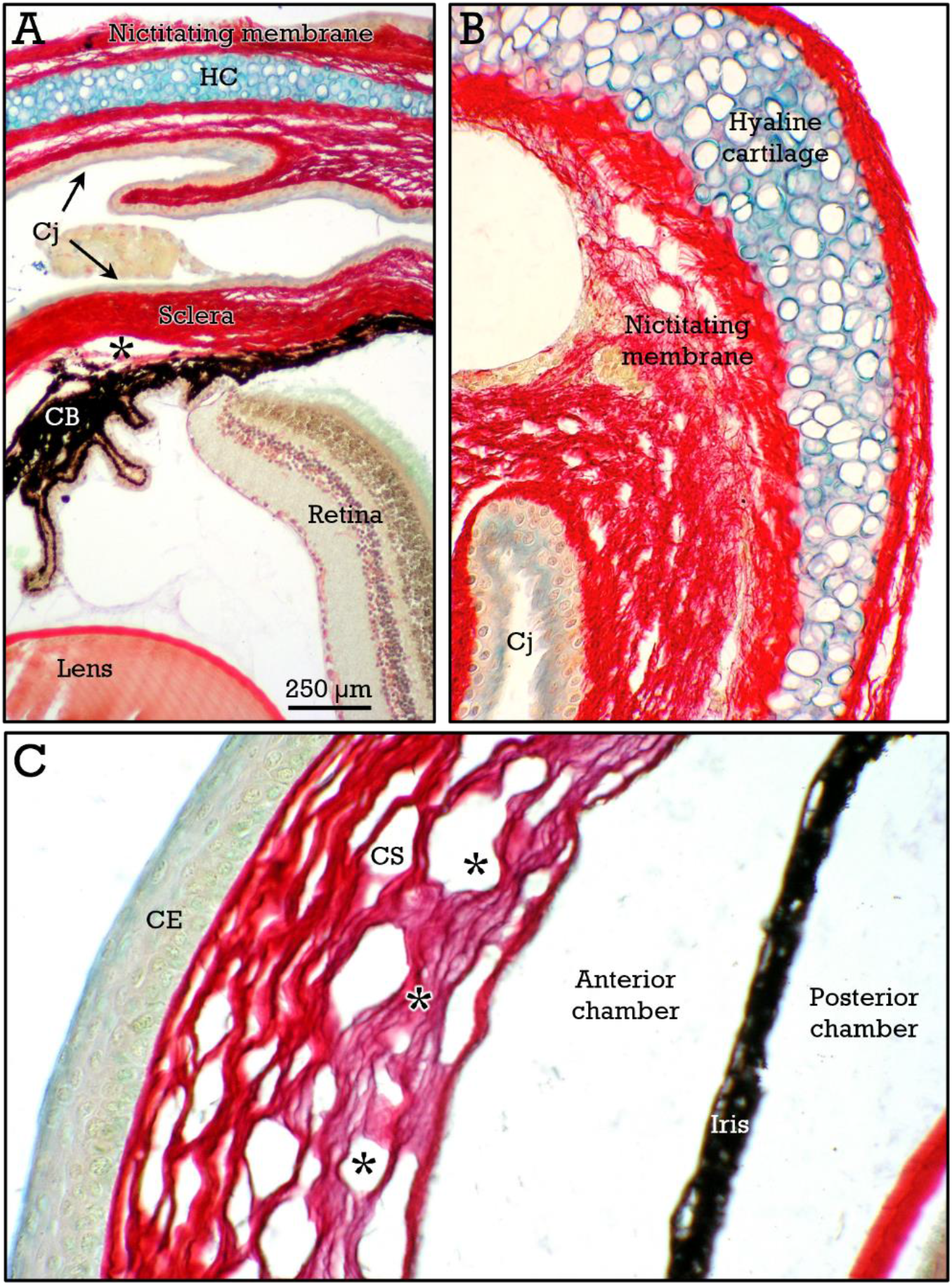
Mouse sclera and cornea. In **A, B**, The conjunctiva (***Cj***) is coating the rudimentary nictitating membrane, that contains a lamina of hyaline cartilage (***HC***). The Schlem canal (***asterisk***) can be observed at the iridocorneal angle. The cornea is composed of the anterior corneal epithelium (***CE***), the stroma (***CS***) and the posterior endothelium (***En***). The outer zone of the corneal stroma stains red, whereas the inner zone stains magenta (***asterisks***). The highly pigmented iris delimitates the anterior and posterior chambers.

### Zebrafish eye

Despite belonging to different vertebrate groups, the general structure of the zebrafish eye, and particularly of the retina, was very similar to that of mice and humans. However, highly significant morphological and functional differences, such as the regenerative capacity of the zebrafish retina, exist.

#### Retina

In the zebrafish, the use of different fixatives results in relevant differences in the staining properties of the retina (Figs. 15A,B). In formaldehyde-fixed samples (Fig. 15B), staining was more similar to that of human and mouse retina. Thus, cell nuclei in the ONL were uniformely red-stained whereas a heterogeneous staining pattern was found in the INL (Fig. 15B). On the other hand, the general layered structure of the zebrafish retina (Fig. 15A,B) was equivalent to that of human and mouse, with a considerably wider retinal pigment epithelium. Notably, tangential sections at the level of the photoreceptor layer show a geometrically precise hexagonal cone mosaic pattern (Fig. 15C). The structure of the optic nerve was also close similar to that of the other species, although abundant melanocytes were present at the *lamina cribosa* (Fig. 16A). Large clusters of melanocytes were also observed surrounding the choroid blood vessels in the proximity of the optic nerve (Fig. 16B).

**Figure 15.**
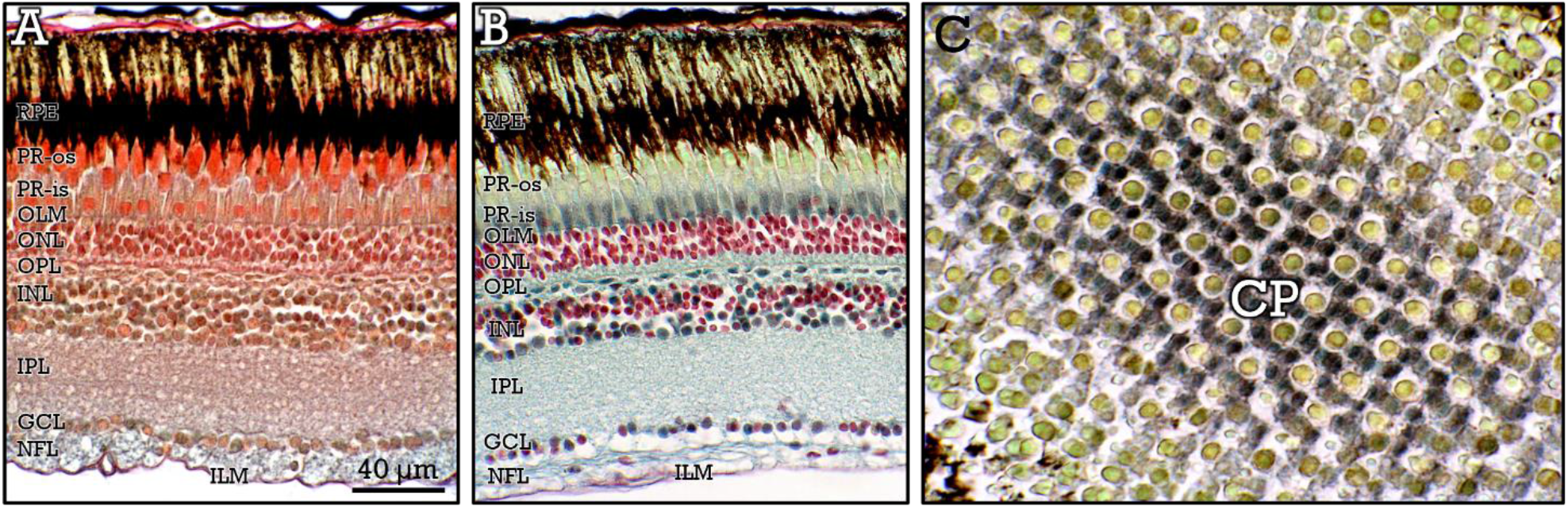
Zebrafish retina. Bouin (**A**) and formaldehyde (**B**) fixed tissues show significant staining differences. The classical retinal layers, that are better defined in formaldehyde fixed samples, are indicated. Tangential sections (**C**) at the level of the inner segment of the photoreceptor layer show the geometrically precise cone pattern.

**Figure 16.**
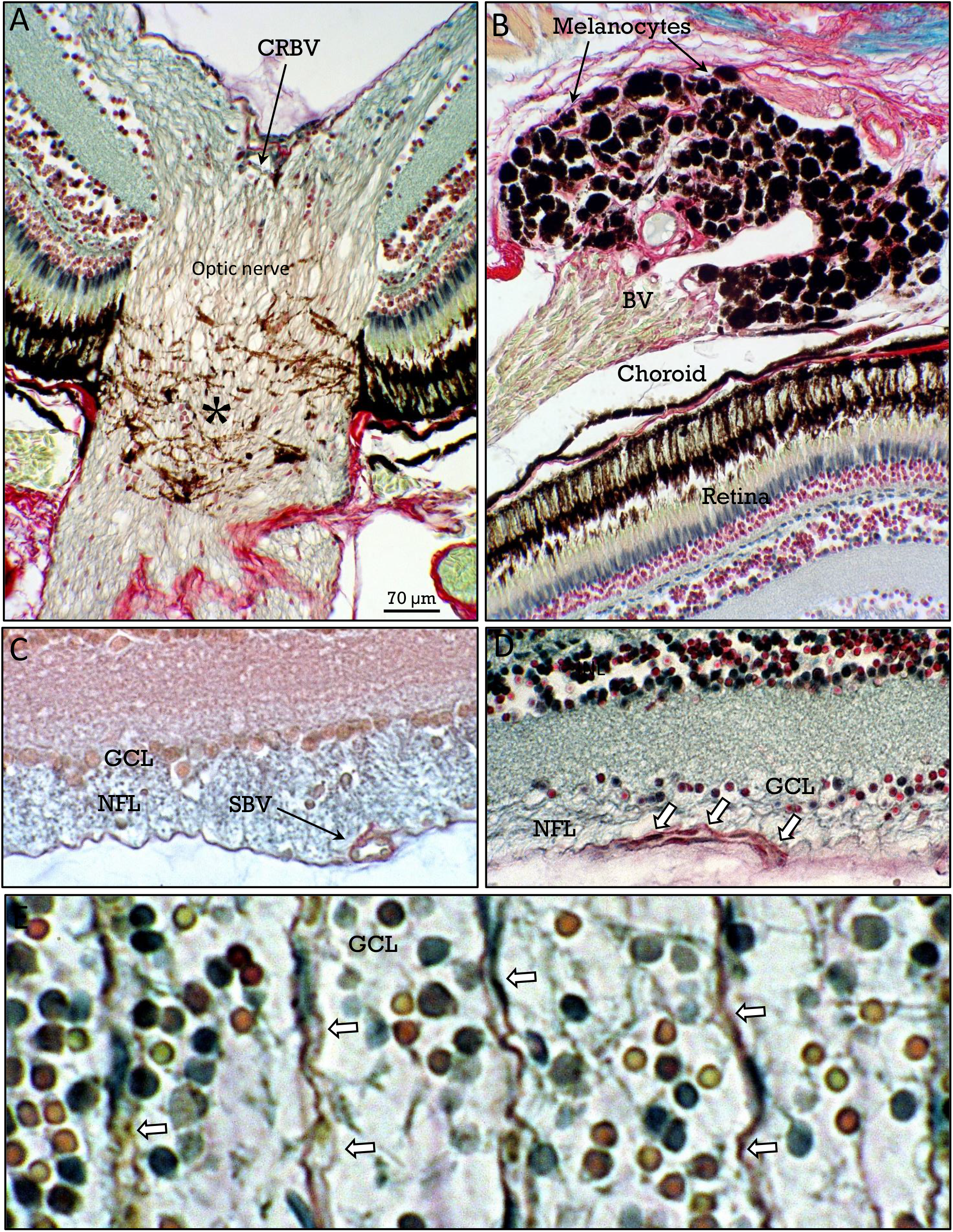
Zebrafish retina vascularization. **A**, central retinal blood vessels (***CRBV***) run along the optic nerve. Melanocytes (***asterisk***) are abundant at lamina cribosa of the optic nerve (**A**) and as large clusters (**B**) adjacent to the blood vessels (***BV***) at the posterior aspect of the globe. Retinal blood vessels form a superficial plexus at the nerve fiber layer (***SBV*** in **C, *arrows* in D**) at the inner retinal surface. A tangential section at the level of the GCL (**E**) shows the heterogeneous staining of cell nuclei, and the abundant underlying capillaries of the superficial plexus (***arrows***).

A dual pattern of vascularization of the retina was also present. Blood vessels entering at the level of the optic nerve, form a superficial blood vessel network at the internal limit of the retina. However, a deeper blood vessel plexus is lacking (Figs. 16C-E). The superficial vascular network was located at the inner surface of the retina (Figs. 16C,D), below the ganglion cell layer that show a clear heterogeneous staining pattern (Fig. 16E). As in other vertebrate species the external retinal layers are nourished by diffusion from the adjacent choroidal vessels.

#### Uvea

As in the other species, the uvea was divided into the choroid, the ciliary body and the iris (Fig. 17A). The choroid contains abundant blood vessels (Fig. 17B) that form a special vascular structure (the *rete mirabilis*) adjacent to the optic nerve (Fig. 17C). The choroid, the ciliary body and the iris were highly melanized (Figs. 17A-E). Except for the fact that the zebrafish lens is spherical, the histological structure and staining properties of the lens were equivalent to that in the other species (Figs. 17F-H).

**Figure 17.**
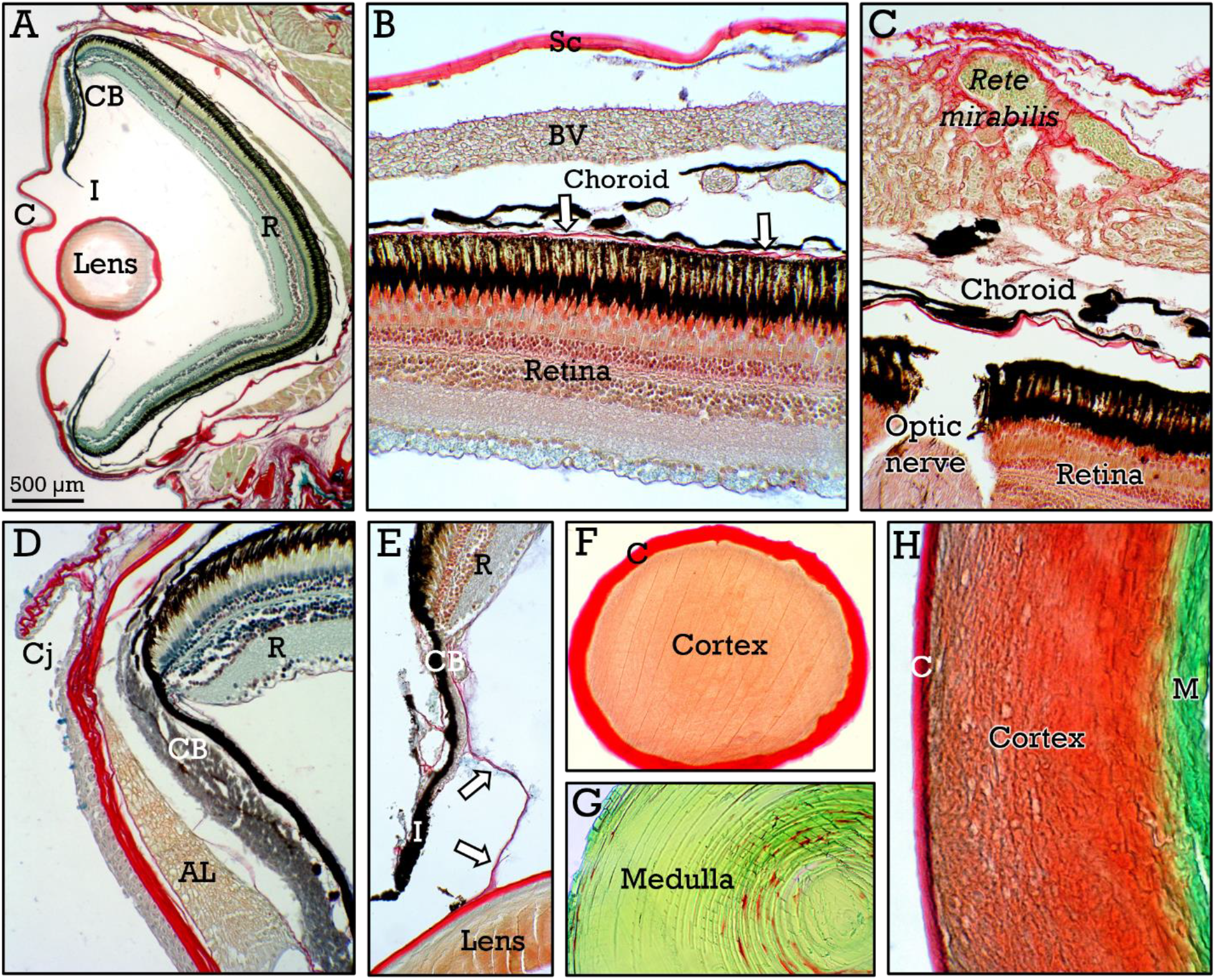
Zebrafish uvea and lens. In **A**, general view of the zebrafish eye. **B**, the choroid contains abundant melanocytes and blood vessels (***BV***) responsible for the irrigation of the external retina. The Bruch membrane (***arrows*** in **B**) is located at the limit between choroid and retina. In **C**, the vascular *rete mirabilis* can be observed at the posterior zone of the choroid. The ciliary body (***CB***), fibers of the zonula (***arrows***) connected to the lens, the iris (***I***), the conjunctiva (***Cj***), the cornea (***C***), and the annular ligament (***AL***) are shown in **D**,**E**. The staining characteristics of the spherical lens were identical to those of human and mouse (**F-H**), showing an intensely red-stained capsule (***C***), a red-stained cortex, and a green-stained medulla (***M***).

#### Sclera and cornea

The sclera appear as a red-stained thin lamina to which extrinsic ocular muscle were inserted (Fig. 18A). At the anterior aspect, it was covered by the conjunctiva and near the limbus it has a cartilaginous ring (Fig. 18B,C). The cornea has an external stratified epithelium, covering a collagenous stroma (Fig. 18D).

**Figure 18.**
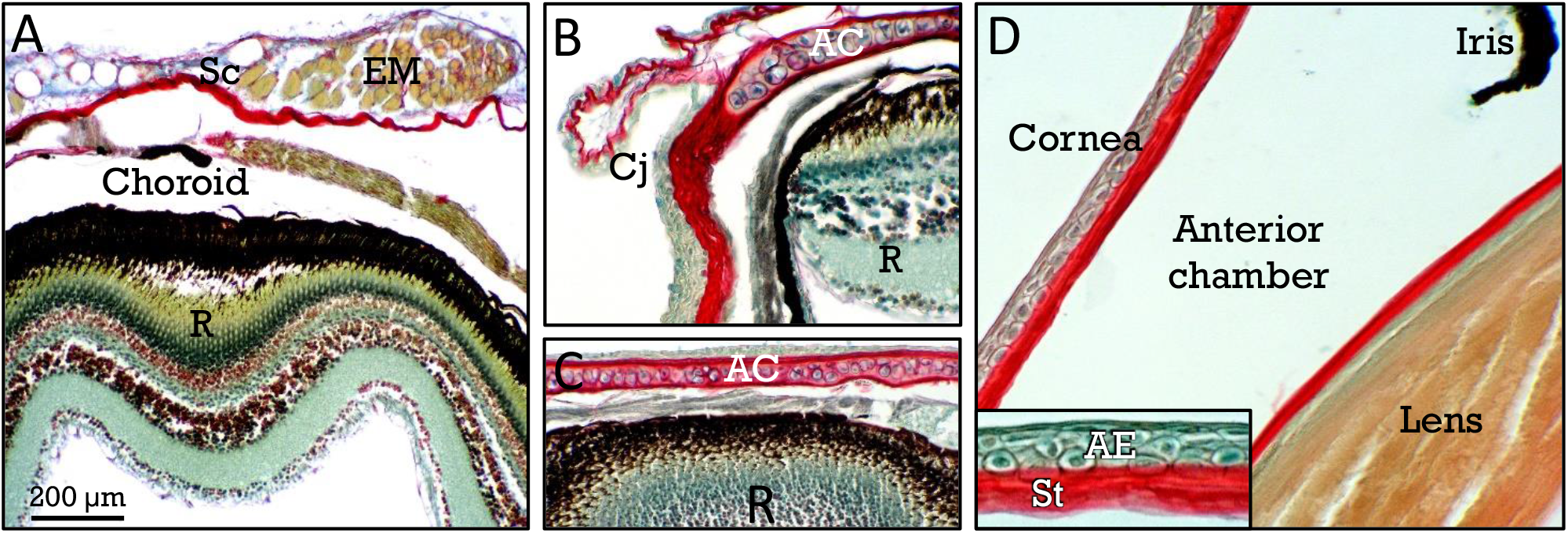
Zebrafish sclera. The sclera appears as a thin red-stained lamina (**A**) to wich extrinsic ocular muscles (***EM***) are inserted. Behind the conjunctiva (***Cj***), the annular cartilage can be observed (***AC*** in **B**,**C**). The cornea show an anterior stratified epithelium (***AE***) and a collagenous stroma (***St***).

## Discusion

The results obtained in this study after the application of the RGB staining method to human, mouse, and zebrafish eye tissues are extensively shown in the figures. These data indicate that RGB trichrome constitutes a simple and reliable staining method, complementary to the routine H&E staining, that results in brilliant tissue staining and highly contrasted tissue interfaces. This facilitates tissue identification and is advantageous for automatic or semiautomatic tissue segmentation and digitization procedures^**15**,**16**^. Consequently, these staining properties are adequate for both eye research and histo(patho)logy teaching. In this setting, the question is to what extent the RGB trichrome actually has any advantage over other widely used trichrome stains when applied to the eye tissues.

The effectiveness of RGB trichrome to distinguish collagens from other tissue components is similar to that of other classical trichrome stains, such as the popular Masson trichrome^**17**,**35**^. However, the RGB trichrome gives additional information about the composition of the extracellular matrix, that plays multiple roles in eye physiology^**36**^. This trichrome stain has been applied to specialized connective tissues such as bone and cartilage^**18**^ resulting in a wide array of shades of color. Two main components of the extracellular matrix of connective tissue proper are collagens and glycosaminoglycans^**37**^, that are selectively stained by picrosirius red and alcian blue respectively. Consequently, the staining color of connective tissues depens on the relative proportion of these two main components. This results in red staining when collagen fibers are predominant, while in connective tissues with a relatively high proportion of glycosaminoglycans color shifts to magenta. This can be clearly appreciated by comparing the stroma of the iris (magenta-stained) with that of ciliary projections (red-stained; Fig. 19 A) or the inner and outer areas of the ciliary body (Fig 19 B) in human eyes, or the mouse sclera with respect to periocular connective tissue (Fig. 19C) or to the stroma of the conjunctiva (Fig. 19D). Also in this sense, different zones of the mouse corneal stroma (anterior half *vs* posterior half) stain differently (Fig. 14C), which is likely related to the uneven distributtion of proteoglycans from front to back of the corneal stroma^**38**^. Finally, in the zebrafish some areas of mucous-like connective tissue near the optic nerve stain blue (Fig. 19E). Summarizing, the combination of sirius red and alcian blue in the RGB stain gives rise to a range of colors, depending on the relative proportions of extracellular matrix components, mainly collagens and glycosaminoglycans/proteoglycans.

**Figure 19.**
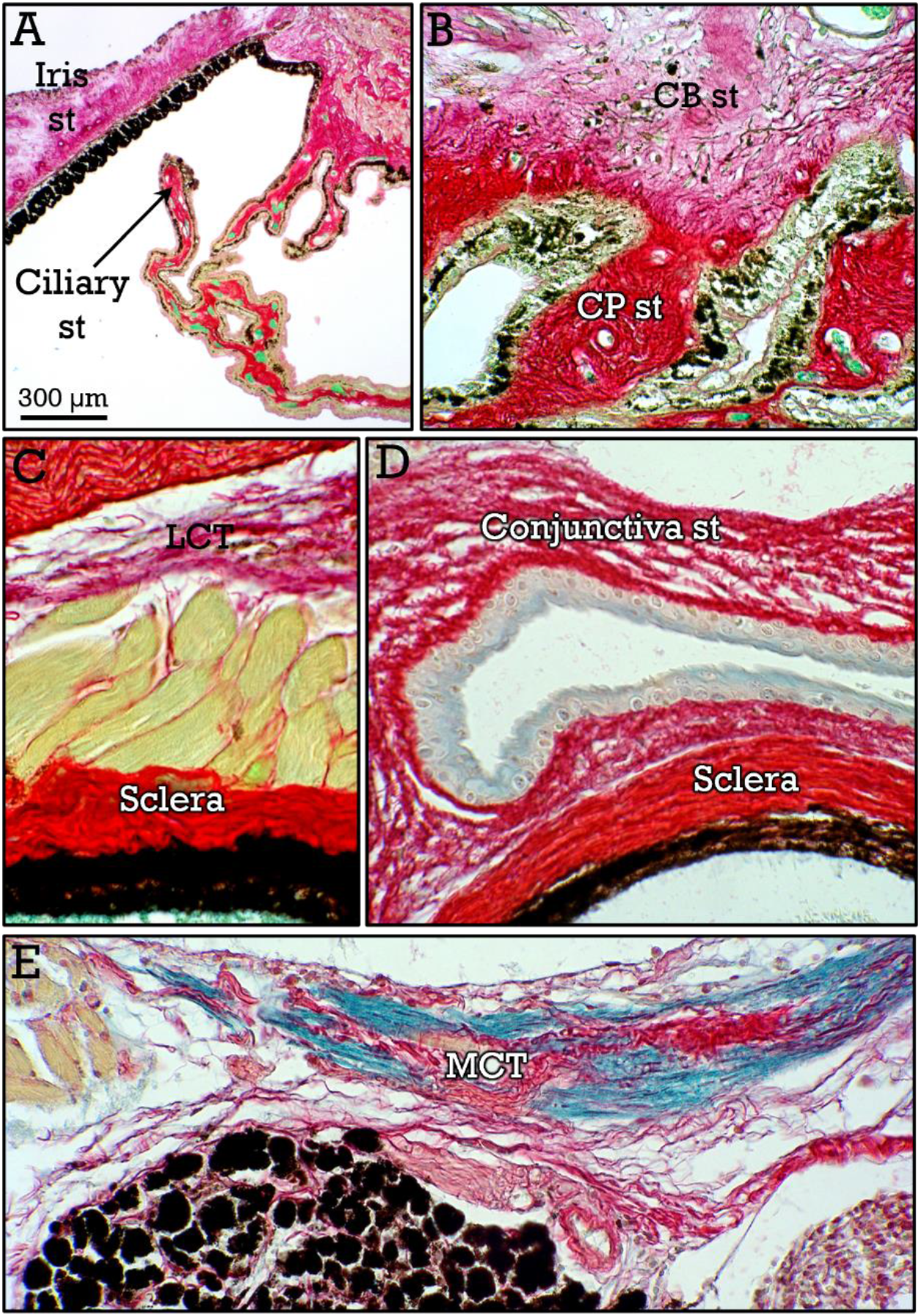
Staining of connective tissue proper. Connective tisssue stains with different shades of colors depending on the proportion of collagen and glycosaminoglycans. Red vs magenta stained connective tissue can be observed at the iris and ciliary body in human samples (**A**,**B**), and sclera *vs* loose connective tissue (***LCT***) and *vs* conjunctiva stroma in the mouse (**C**,**D**). In the zebrafish, blue stained areas of mucous connective tissue (***MCT***) can be observed near the optic nerve (**E**).

Staining of the retina – that is the most remarkable eye tissue – with RGB trichrome is particularly relevant, when compared to that of other conventional methods. Notably, the application of RGB stain gives rise to a differential staining of the retinal layers. So, the photoreceptor outer segments stain blue, whereas inner segments stain green. The blue staining of the outer photoreceptor segments seems to be due to the presence of some specific proteoglycans in the interphotoreceptor matrix^**39**,**40**^.

Furthermore, the outer and inner nuclear layers show different staining patterns, showing the ONL a homogeneous staining, in contrast to the INL that was heterogeneously stained. The heterogeneous staining of the INL is likely due to the presence of the nuclei of different cell types, such as bipolar, amacrine, and horizontal neurons (and their subtypes), as well as Müller glial cells in this layer^**25**,**29**,**32**^. In good agreement, these staining patterns of the ONL and INL were observed in the three species analyzed, although with different species-specific shades of color in the ONL.

In addition, heterogeneous staining was also found in the GCL, that was particularly evident in the zebrafish retina. In this layer, several cell types such as astrocytes and microglial cells are present, in addition to several subtypes of ganglion cell neurons^**31**,**41**^. It is tempting to speculate that differently stained nuclei in both INL and GCL layers correspond to different specific cell types. However, matching staining colors with specific cell types was far beyond the purposes of this study.

In summary, the RGB trichrome is effective for the differentiation of eye tissues, producing highly contrasted tissue interfaces. Furthermore, staining of the retina, the organ-specific tissue in the eye, yields to a differential staining of the neuronal retinal layers. These properties make RGB stain a reliable tool for ophthalmologic studies in human pathology and basic preclinical research, as well as for histo(patho)logy teaching.

## Acknowledgements

The authors are very grateful to Dr M Barasona and Dr MT Urbano from the Servicio de Animales de Experimentación of the University of Cordoba for the zebrafish samples, and Dr E Tarradas for his technical support.

## Author contributions

Both authors were involved in conducting the study and contributed to the final version of the manuscript.

